# Multifunctional Roles of Sec13 Paralogues in the Euglenozoan *Trypanosoma brucei*

**DOI:** 10.1101/2024.12.03.626618

**Authors:** Mohamed Sharif, Lydia Greenberg, James Bangs

## Abstract

Secretory cargos are exported from the ER via COPII coated vesicles that have an inner matrix of Sec23/Sec24 heterotetramers and an outer cage of Sec13/Sec31 heterotetramers. In addition to COPII, Sec13 is part of the nuclear pore complex (NPC) and the regulatory SEA/GATOR complex in eukaryotes, which typically have one Sec13 orthologue. The kinetoplastid parasite *Trypanosoma brucei* has two paralogues: TbSec13.1, an accepted component of both COPII and the NPC, and TbSec13.2. Little is known about TbSec13.2, but others have proposed that it, and its orthologue in the distantly related diplonemid *Paradiplonema papillatum*, operate exclusively in the SEA/GATOR complex, and that this represents an evolutionary diversification of function unique to the euglenozoan protists (doi.org/10.1098/rsob.220364). Using RNAi silencing in trypanosomes we show both TbSec13s are essential. Knockdown of each dramatically and equally delays transport of GPI-anchored secretory cargo, indicating roles for both in COPII-mediated trafficking from the ER. Immunofluorescence and proximity labeling studies confirm that both TbSec13.1 and TbSec13.2 co-localize with TbSec24.1 to ER exit sites, and thus are functional components of the COPII machinery. Our findings indicate that TbSec13.2 function is not restricted to the SEA/GATOR complex in trypanosomes.

## INTRODUCTION

The protozoan parasite *Trypanosoma brucei* (*T. brucei ssp*) is the causative agent of Human African Trypanosomiasis (HAT, Sleeping Sickness) and the etiologically and epidemiologically similar African Animal Trypanosomiasis (AAT, Nagana) in domestic livestock [1, 2]. The parasite has a dixenous life cycle alternating between a mammalian host and an insect vector, the tsetse fly [3]. HAT and AAT are endemic in 36 sub-Saharan African countries with tsetse flies (World Health Organization, www.who.int). A major virulence factor in disease progression within the mammalian host is the expression of antigenically distinct variant surface glycoproteins (VSG) by bloodstream form parasites (BSF) [4]. Densely packed on the cell surface, VSG shields underlying invariant surface proteins from host immune recognition [5–7]. Only one VSG variant is expressed at any given time, and the process of switching VSG is known as antigenic variation. VSG is the major glycosylphosphatidylinositol (GPI)-anchored protein in trypanosomes, and while other GPI-anchored proteins exist, VSG accounts for approximately 10% of the total protein synthesized in BSF trypanosomes [8], and thus the overwhelming amount of all secretory cargo. To accommodate this need, trypanosomes have evolved a highly efficient and streamlined secretory pathway [9, 10].

Like all secretory cargoes in eukaryotes, VSG is synthesized in the endoplasmic reticulum (ER), where it is N-glycosylated and GPI-anchored prior to export via ER exit sites (ERES) [9, 11–14]. At the ERES, cargo is packaged into COPII coated vesicles for transport to the downstream Golgi apparatus [15, 16]. Coat assembly is initiated by deposition of activated GTP-bound Sar1 at ERES budding sites resulting in recruitment of Sec23/Sec24 heterodimers that form the inner COPII layer. It is this ‘pre-budding’ complex that is responsible for cargo recruitment to budding vesicles. Subsequent recruitment of outer Sec13/Sec31 heterotetramers leads to membrane deformation and vesicle scission. Trypanosomes have orthologues of all the main COPII coat components (Table 1), including two paralogues each of TbSec23, TbSec24 and TbSec13. Our previous work demonstrated that the TbSec23/TbSec24 subunits form specific and obligate heterodimers: Pair A (TbSec23.2/TbSec24.1) and Pair B (TbSec23.1/TbSec24.2) [9]. In BSF trypanosomes, GPI anchors are forward trafficking signals for ER exit [14, 17, 18]. Deletion of the GPI attachment peptide from VSG delays transport resulting in accumulation in the ER, and attachment of a GPI peptide to soluble reporters can accelerate exit. This GPI-dependent transport is specifically mediated by the Pair A Sec23/Sec24 heterodimer, in conjunction with transmembrane adaptors (TbERPs) that recognize GPI in the lumen and TbSec24.1 in the pre-budding complex [9, 14].

**Table 1.** *T. brucei* & *P. papillatum* COPII orthologues

| Subunit | *T. brucei*<sup>1</sup> | *P. papillatum*<sup>2</sup> |
| --- | --- | --- |
| **Sar1**<sup>3</sup> | Tb927.5.4500 | DIPPA_22352 |
| **Sec23.1** | Tb927.8.3660 | DIPPA_30134 |
| **Sec23.2** | Tb927.10.7740 | DIPPA_35143 |
| **Sec24.1** | Tb927.3.1210 | DIPPA_32844 |
| **Sec24.2** | Tb927.3.5420 | DIPPA_07091 |
| **Sec13.1** | Tb927.10.14180 | DIPPA_24951 |
| **Sec13.2** | Tb927.11.8120 | DIPPA_35934 |
| **Sec31** | Tb11.02.4040 | DIPPA_09941 |

<sup>1</sup> Sevova and Bangs (2009) *Mol. Biol. Cell* 20:4739
<sup>2</sup> Faktorova et al. (2023) *Open Biol.* 13:220364
<sup>3</sup> *P. papillatum* also has a Sar1B orthologue, DIPPA\_03493

In the Sec31:Sec13 heterotetramer Sec13 binds to a flexible region in the N-terminal half of each Sec31 subunit, between a β-propeller and an α-solenoid domain [16, 19]. This provides rigidity to the overall structure, which in turn facilitates membrane deformation as the COPII cage is assembled. Sec13 also functions in other cellular processes. It is a widely conserved structural component of the nuclear pore complex (NPC) outer ring, including in trypanosomes [20, 21], and it is part of the likewise broadly conserved SEA/GATOR complex, an essential regulator of the mTORC1 sensing pathway with localization to lysosomal/vacuolar membranes [22]. Yeast and mammals each have a single copy of Sec13, which participates in all of these functions. However, trypanosomes have two paralogues, TbSec13.1 and TbSec13.2, as does the distantly related euglenozoan (diplonemid) *Paradiplonema papillatum* (Table 1). TbSec13.1 is a *bona fide* component of the trypanosome NPC [20, 21], and has been localized to the ERES consistent with a role in COPII vesicles [23]. The TbSec13.2 orthologue was not investigated in these studies. The *P. papillatum* orthologues, PpSec13a and PpSec13b respectively, have been studied [24], and based on localization and pull-down proteomic analyses it was concluded that PpSec13a has dual function in the NPC and in COPII vesicles, but that PpSec13b is solely involved in SEA/GATOR function. By analogy this dichotomy was extended to the *T. brucei* orthologues, although no functional studies were performed in either species.

In this work we perform immunofluorescent and proximity labeling localization studies in trypanosomes, and use an RNAi knock down approach to assess the role of both TbSec13.1 and TbSec13.2 in secretory trafficking from the ER, i.e, COPII function. Our rationale is two-fold. First, we wish to know if the subunit specificity of GPI-dependent ER exit seen with the Sec23/Sec24 heterodimer in the inner COPII coat extends to the two Sec13 subunits in the outer coat. Second, we wish to test the strict functional dichotomy proposed by Faktorova et al. [24] for the two Sec13 orthologues in euglenozoan protozoa. Our results provide definitive answers to both these questions in *T. brucei*.

## RESULTS

### Identification of trypanosomal Sec13 paralogues

We previously identified two paralogous trypanosomal Sec13 genes by querying the TriTryp genomic database (https://tritrypdb.org/tritrypdb/app) with the *Saccharomyces* orthologue (YLR208W) [9]. These were denoted as TbSec13.1 (Tb927.10.14180) and TbSec13.2 (Tb927.11.8120). A recent study in the distantly related and free living marine euglenozoan *Paradiplonema papillatum* referred to these paralogues as Sec13a and Sec13b, respectively [24]. We will adhere to our original designation throughout this report so as to conform to our long established nomenclature for trypanosomal COPII subunits (Table 1) [9, 14, 25, 26].

### TbSec13.1 and TbSec13.2 are essential in BSF trypanosomes

Our previous knockdown studies of the inner COPII coat components (Pair A: TbSec23.2/TbSec24.1; Pair B: TbSec23.1/TbSec24.2) indicated that both heterodimers are essential in BSF cells [9]. In each case the transport of transmembrane (p67) or soluble (TbCatL) cargoes were largely unaffected, suggesting functional redundancy. However, transport of GPI-anchored cargo was uniquely dependent on Pair A. While it is unknown if Pair A forms a distinct homotypic class of COPII vesicles, or whether there is a single heterotypic class containing both Pair A and B, these initial findings raise the question of whether this GPI-selective transport extends to the COPII outer layer. We now investigate the role of TbSec13 paralogues in GPI-dependent trafficking using conditional RNAi constructs independently targeting either the TbSec13.1 or TbSec13.2 subunit. In both cases, RNAi silencing in BSF trypanosomes resulted in the loss of cell viability (Fig. 1). For TbSec13.1, sustained growth arrest was observed as early as 12 hr, and complete cell death occurred at 24 hr (Fig. 1A, left). Knockdown efficiency was assessed at 8 hrs, when cell morphology appeared normal (data not shown), using quantitative real-time PCR (qRT-PCR). Silencing specifically reduced TbSec13.1 transcript levels to 43.7 ± 0.1% (mean ± SD, n = 3) without affecting TbSec13.2 message levels (Fig. 1A, right). For TbSec13.2, sustained growth arrest was observed after 24 hr, and complete cell death occurred by 36 hr (Fig. 1B, left). Knockdown efficiency was assessed at 18 hr, when cell morphology appeared normal (data not shown). Silencing specifically reduced TbSec13.2 transcript levels to 56.7 ± 0.1% (mean ± SD, n = 3) without affecting TbSec13.1 message levels (Fig. 1B, right). Collectively, these data indicate that both TbSec13 subunits are critical for cell viability in BSF trypanosomes.

**Figure 1.**
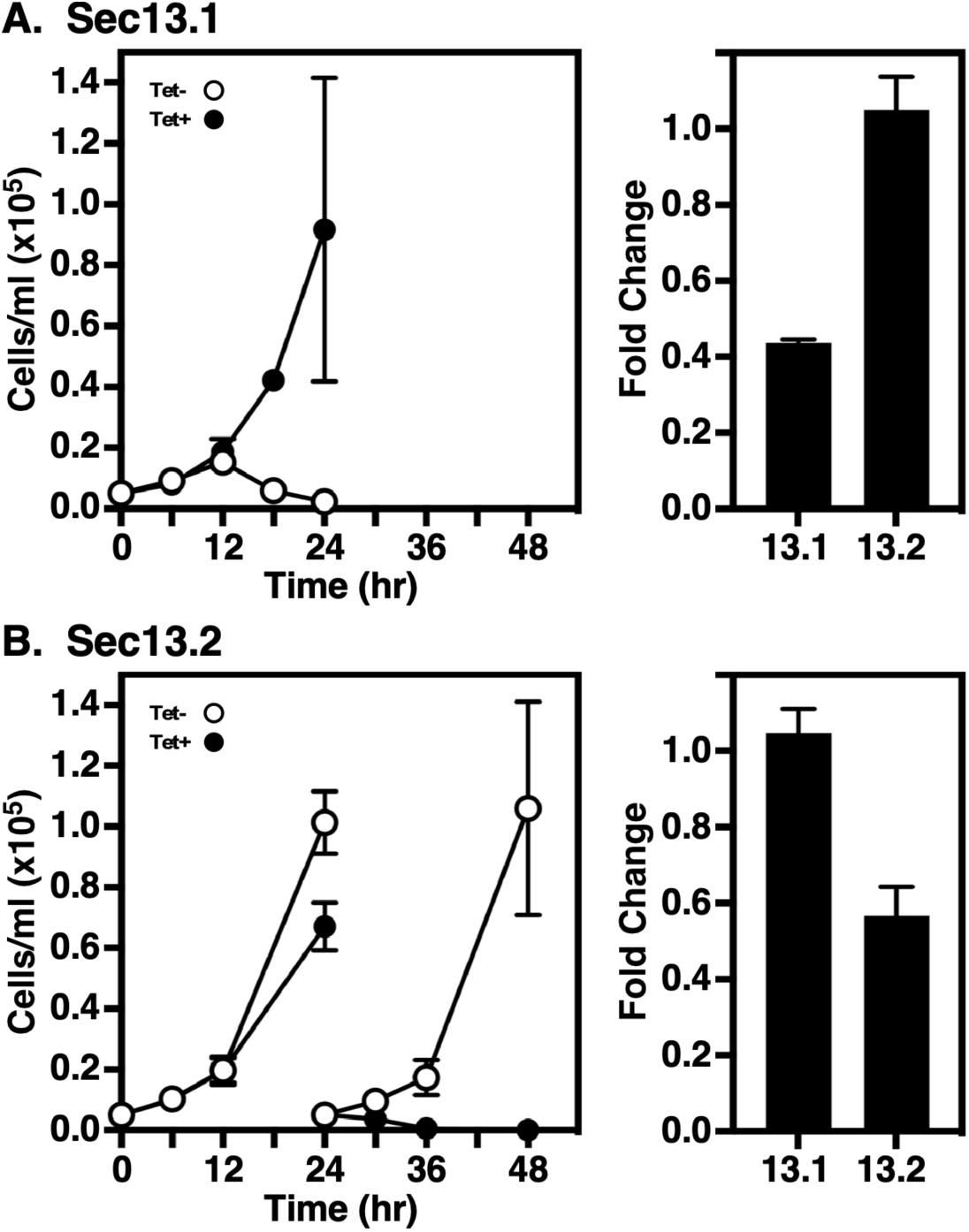
Silencing TbSec13.1 or TbSec13.2 subunits. TbSec13.1 (A) and TbSec13.2 (B) RNAi cell lines were cultured without (open circles) and with (closed circles) tetracycline to initiate dsRNA synthesis. **Left.** Cells were seeded at 5×10^4^ cells/ml and counted every 6 hrs. After 24 hrs, cells were adjusted to the starting density to maintain log phase growth. **Right.** mRNA levels in control (Tet-) and silenced (Tet+) cells were determined by qRT-PCR at 8 hrs (TbSec13.1) and 18 hrs (TbSec13.2) of induction. mRNA levels were normalized using the internal control, ZFP3. Data are presented as the fold change from uninduced control. All growth and qRT-PCR assays were performed in triplicate, and three biological replicates were conducted. The data are presented as mean ± SD.

### Both TbSec13 paralogues are required for efficient ER exit of GPI-APs

The presence of two TbSec13 paralogues raises the question of whether they are functionally redundant, or whether they play distinct cargo-specific roles with regards to ER exit of secretory cargo. To investigate this, we first analyzed the trafficking of endogenous GPI-anchored VSG221 after specific RNAi silencing. Pulse/chase radiolabeling was performed, and arrival of VSG at the cell surface was quantified by the hypotonic lysis assay [9, 11]. Upon arrival at the cell surface, VSG is susceptible to release by the action of endogenous GPI-PLC after hypotonic lysis, while internal VSG *en route* to the surface is resistant. Knockdown of either TbSec13.1 or TbSec13.2 subunits delayed VSG transport from the ER to the cell surface (Fig. 2). Precise half-times (*t*_1/2_s) determined by nonlinear regression are presented in Table 2. For TbSec13.1 and TbSec13.2 cell lines, the calculated VSG transport half-times under normal conditions were 0.18 hr (10.8 min) and 0.12 hr (7.2 mins), respectively. In each case, knockdown resulted in in a 3-to 4-fold delay in VSG transport. This delay in VSG transport is statistically significant (p-value ≤ 0.05) as indicated by non-overlapping 95% Confidence Intervals (CI) ranges for each data set (Table I). These data indicate that both TbSec13 subunits are required collectively for efficient GPI-dependent ER exit.

**Figure 2.**
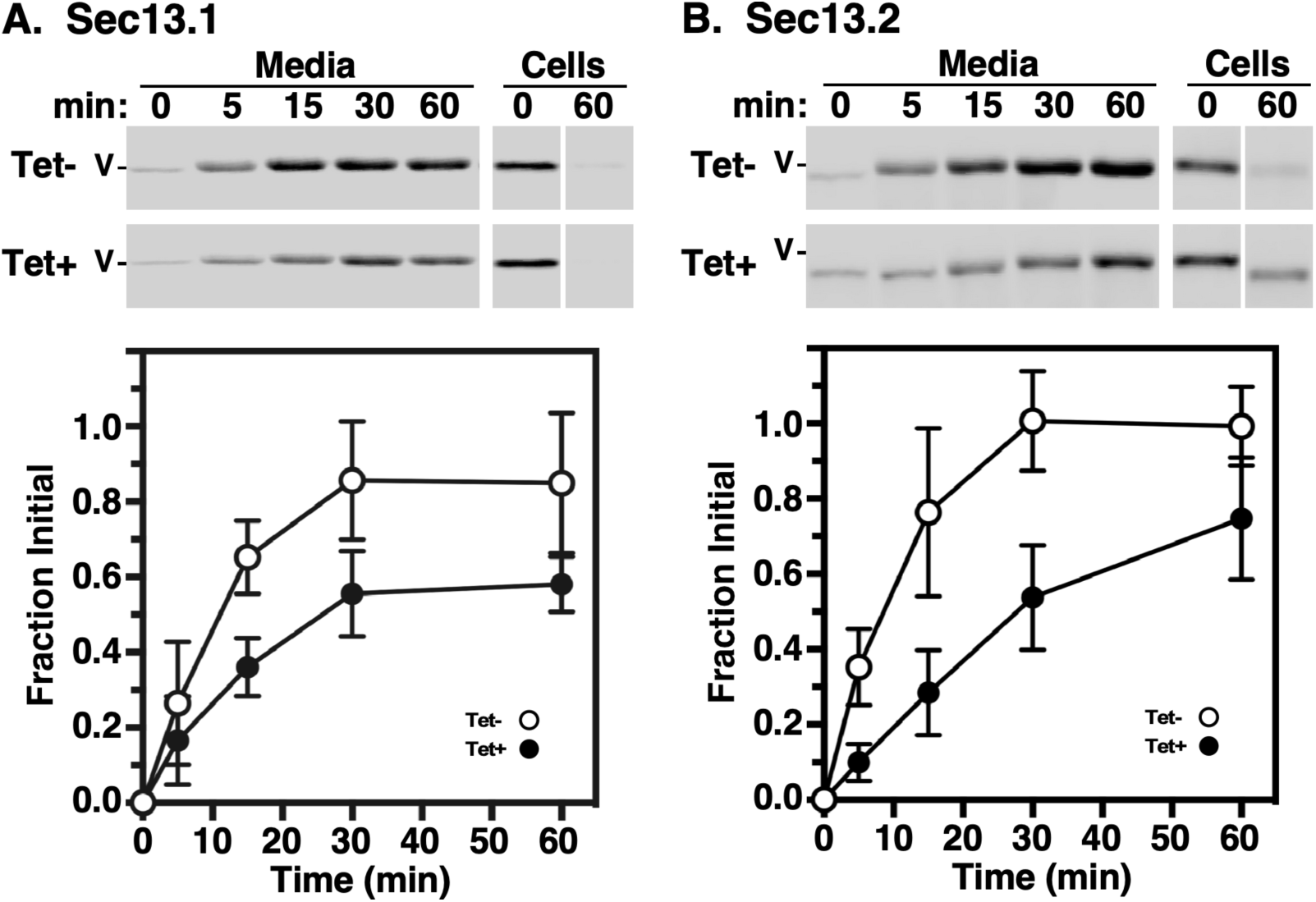
VSG transport in TbSec13 Knockdowns. Specific dsRNA synthesis was induced in the TbSec13.1 (A, 8 hr) and TbSec13.2 (B, 18 hr) RNAi cell lines, and transport of newly synthesized VSG to the cell surface was assessed by the hypotonic release procedure (see Methods). Cells were pulse (2 min)/chase (60 min) radiolabeled and released fractions were prepared by centrifugation at the indicated chase times. VSG221 polypeptides were specifically immunoprecipitated and analyzed by SDS-PAGE/phosphorimaging. **Top.** Representative images for control (Tet-) and silenced (Tet+) cells are presented (10^6^ cell equivalents per lane). All vertical white spaces indicate lanes that were excised post-image processing for the sake of presentation. Matched Tet– and Tet+ gels are from the same processed phosphorimage. Mobility of VSG (V) is indicated. **Bottom.** Quantification of the released fraction indicating arrival at the cell surface. All values are normalized to T_0_ total (mean ± std. dev., n=3 biological replicates).

**Table 2.** Kinetics of Reporter Transport^a^

| Reporter | RNAi Target | Tet | $t_{1/2}$ (hr) | 95% CI (hr)<sup>b</sup> | R<sup>2</sup> |
| --- | --- | --- | --- | --- | --- |
| **VSG<sup>c</sup>** | TbSec13.1 | - | 0.18 | 0.13-0.25 | 0.87 |
|  |  | + | 0.53 | 0.40-0.71 | 0.77 |
|  | TbSec13.2 | - | 0.12 | 0.09-0.16 | 0.92 |
|  |  | + | 0.49 | 0.39-0.60 | 0.90 |
| **TbCatL<sup>d</sup>** | TbSec13.1 | - | 0.14 | 0.12-0.17 | 0.96 |
|  |  | + | 0.15 | 0.13-0.17 | 0.97 |
|  | TbSec13.2 | - | 0.16 | 0.14-0.19 | 0.95 |
|  |  | + | 0.18 | 0.16-0.20 | 0.96 |
| **p67<sup>e</sup>** | TbSec13.1 | - | 0.51 | 0.44-0.58 | 0.98 |
|  |  | + | 1.44 | 1.13-1.82 | 0.86 |
|  | TbSec13.2 | - | 0.64 | 0.49-0.82 | 0.90 |
|  |  | + | 1.14 | 0.94-1.38 | 0.92 |

a. Halftimes and 95% Confidence Intervals (CI) were calculated by non-linear regression (see Fig. S1), and the half-times are presented in hrs.
b. By definition, in comparing matched Tet-/± data sets, any non-overlap in 95% CI ranges have P-values of ≤0.05 [44, 45].
c. Measured as loss of full-length VSG from cell fraction.
d. Measured as loss of initial precursors (X + I).
e. Measured as loss of initial gp100 ER glycoform.

### TbSec13 subunits are functionally redundant in TbCatL transport

Next we analyzed the trafficking of soluble secretory cargo using cathepsin L (TbCatL), an endogenous soluble lysosomal hydrolase as a reporter [27]. In the ER, TbCatL is synthesized as 53 (I) and 50 kDa (X) proproteins. These precursors are transported to the lysosome for proteolytic processing resulting in a single active mature form (M, 44 kDa). To determine the roles of TbSec13 subunits in TbCatL trafficking from the ER to the lysosomes, we quantified the loss of initial precursors (I+X) upon arrival in the lysosome. Independent knockdown of TbSec13 subunits did not affect TbCatL ER exit (Fig. 3). For the TbSec13.1 and TbSec13.2 cell lines, the measured transport rates under normal conditions were *t*_1/2_ 0.14 hrs (8.6 mins) and *t*_1/2_ 0.16 hrs (9.79 mins), respectively (Table 2). Specific silencing had no significant effect on these transport rates, suggesting that TbSec13.1 and TbSec13.2 subunits are functionally redundant for TbCatL trafficking.

**Figure 3.**
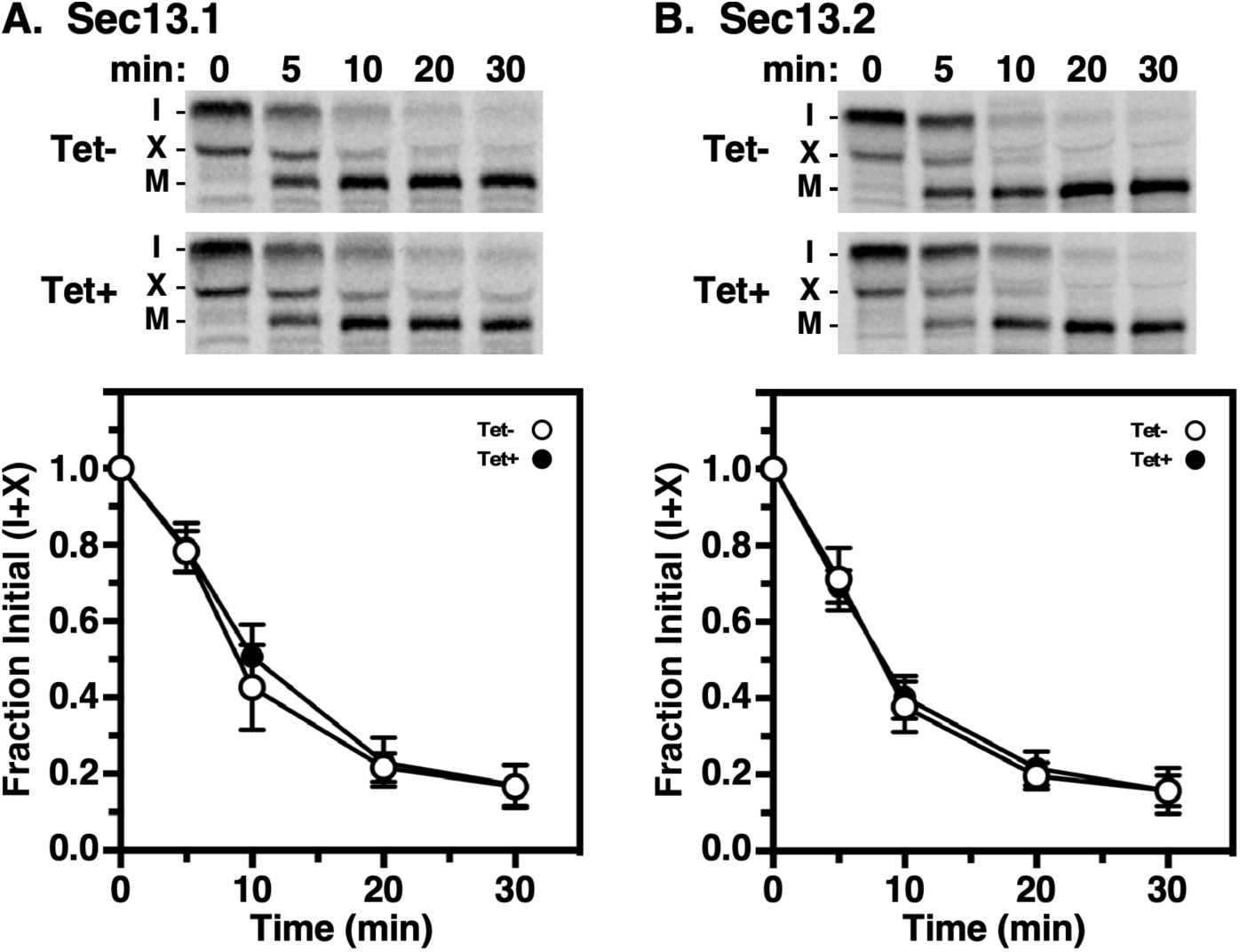
Transport of TbCatL in TbSec13 Knockdowns. Specific dsRNA synthesis was induced for 8 hrs in TbSec13.1 (A) and for 18 hrs in TbSec13.2 (B) RNAi cell lines and pulse (10 min)/chase (30 mins) radiolabeling was performed. TbCatL was immunoprecipitated from cell lysates at the indicated chase times and analyzed by SDS-PAGE and phosphorimaging (10^7^ cells/lane). **Top.** Phosphorimages of representative matched gels from control (Tet −; upper) and silenced (Tet +; lower). Mobilities of initial precursors (I and X) and the lysosomal mature (M) form are indicated. Matched Tet- and Tet+ gels are from the same processed phosphorimage. **Bottom.** Quantification of loss of the initial precursors (I and X). Three biological replicates are quantified, and the data are presented as mean ± SD.

### Both TbSec13 paralogues are required for efficient p67 transport

Finally, we examined the trafficking of p67, a lysosomal-associated type I membrane glycoprotein [9, 28]. In BSF trypanosomes, p67 is initially synthesized in the ER as 100 kDa N-glycosylated protein (gp100). Subsequent N-glycan modification in the Golgi converts gp100 to a 150 kDa glycoform (gp150). From the Golgi, it is transported to the lysosome, where proteolytic fragmentation generates smaller quasi-stable 42 kDa and 32 KDa glycoforms. To determine the roles of TbSec13 subunits in p67 trafficking from the ER, we quantified the loss of gp100 upon transport to the Golgi. Knockdown of either TbSec13 subunits resulted in delays in ER exit (Fig. 4). For the TbSec13.1 and TbSec13.2 cell lines, the calculated ER exit rates under normal conditions were *t*_1/2_ 0.51 hr (30.6 min) and *t*_1/2_ 0.64 hr (38.4 min), respectively (Table 2). Silencing TbSec13.1 or TbSec13.2 subunits resulted in statistically significant delays of 2.8- and 1.8-fold, respectively (Table I). These data support a model in which both TbSec13 subunits are required for efficient ER exit of p67, even though TbSec13.1 silencing had a greater delay than TbSec13.2.

**Figure 4.**
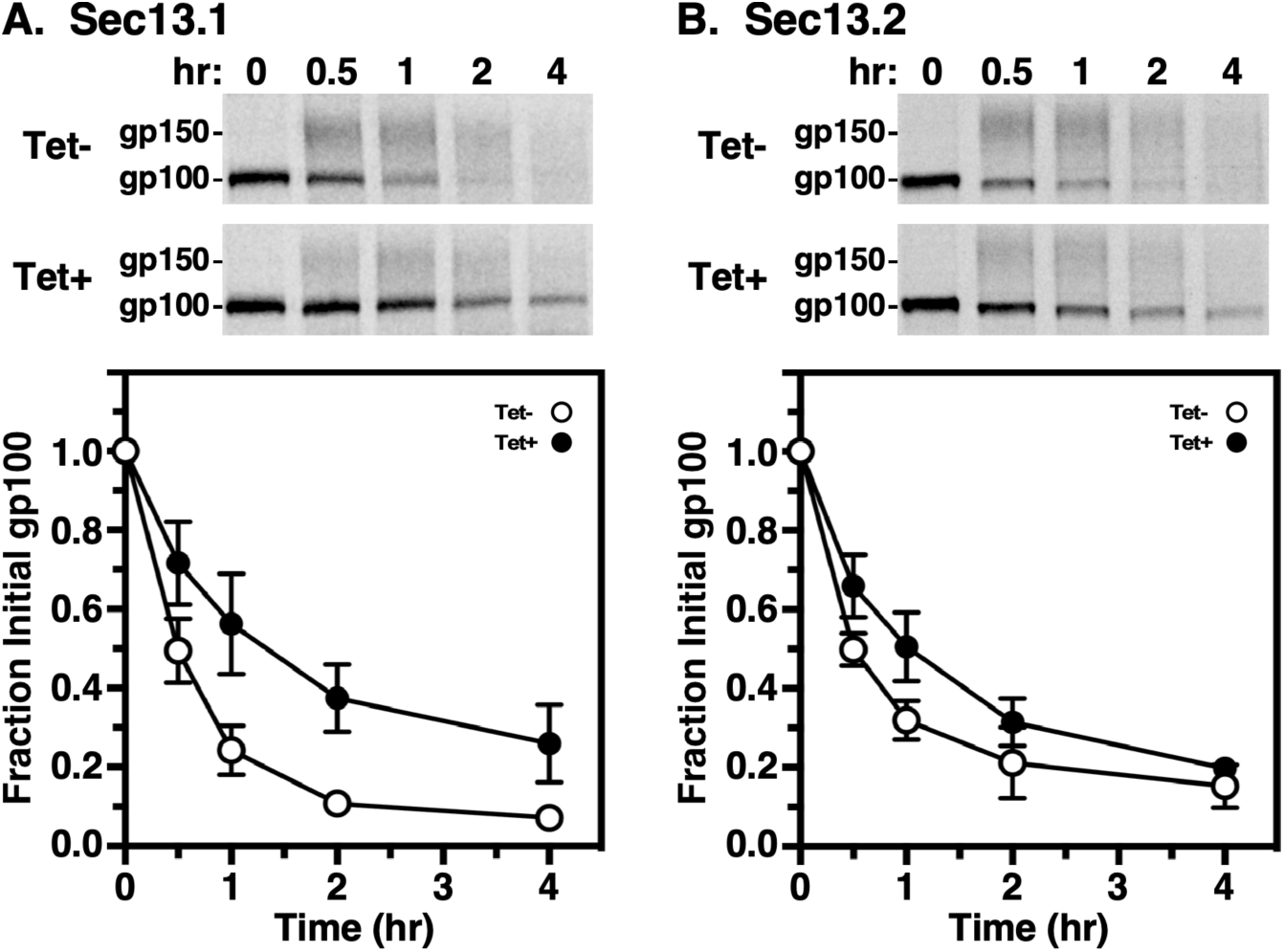
p67 transport in TbSec13 Knockdowns. Specific dsRNA synthesis was induced for 8 hrs in TbSec13.1 (A) and for 18 hrs in TbSec13.2 (B) RNAi cell lines, and pulse (15 min)/chase (4 hrs) radiolabeling was performed. p67 was immunoprecipitated from cell lysates at the indicated chase times and analyzed by SDS-PAGE and phosphorimaging (10^7^ cells/lane). **Top.** Phosphorimages of representative matched gels from control (Tet-) and silenced (Tet+). Mobility of the gp100 precursor and gp150 processed glycoforms are indicated. Matched Tet- and Tet+ gels are from the same processed phosphorimage. **Bottom.** Quantification of loss of the initial ER precursor gp100. Three biological replicates are quantified, and the data are presented as mean ± SD.

### Localization of TbSec13s

For localization studies, both TbSec13 paralogues were independently HA-epitope tagged by *in situ* chromosomal recombination in a BSF host cell line that has a Ty-tagged TbSec24.1 allele as an ERES marker [9]. Western blot analysis (Fig. 5) confirmed proper tagging of TbSec24.1 (108 kDa, top, lanes 2-4), TbSec13.1 (42 kDa, bottom, lane 3) and TbSec13.2 (35 kDa, bottom, lane 4). Interphase (1 kinetoplast, 1 nuclei) BSF cells typically have 2 ERES in the post-nuclear region closely aligned with the extracellular flagellum [25]. Immunofluorescent staining of the TbSec13.1::HA cell line revealed two prominent extra-nuclear spots that co-localized with TbSec24.1::Ty (Fig. 6A) and well-aligned to the flagellum. No obvious staining of the nuclear envelop was observed, in contrast to the published findings of DeGrasse et al. [20] (discussed below). Likewise, TbSec13.2::HA co-localized precisely with TbSec24.1::Ty and in alignment with the flagellum, clearly demonstrating for the first time that it is part of the ERES COPII machinery (Fig. 6B). No other obvious staining of the post-nuclear endolysosomal region that could be construed as indicating association with the SEA/GATOR nutritional complex [24] was observed (discussed below).

**Figure 5.**
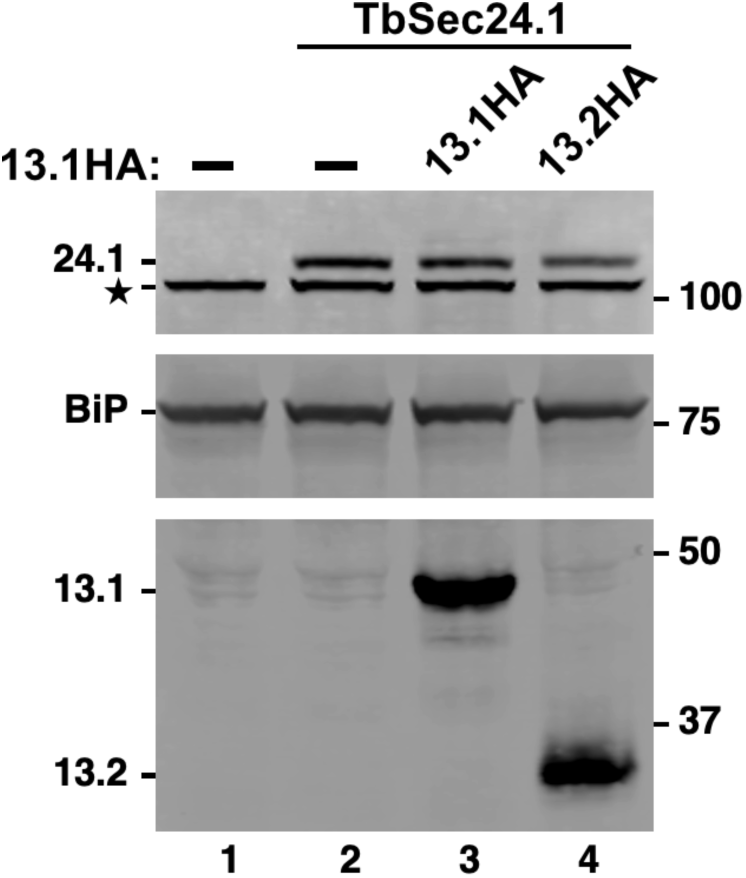
TbSec13 HA-Tagging. Control untagged cells (lane 1) and cells bearing a Ty tagged allele of *TbSec24.1* without (lane 2) or with HA-tagged alleles of *TbSec13*s (lanes 3 & 4) were fractionated by SDS-PAGE, transferred to membranes and probed simultaneously with mAb anti-Ty (top), rabbit anti-BiP (middle) and rabbit anti-HA (bottom). Blots were developed with appropriate secondary reagents and the image was separated digitally for presentation. Mobilities of all targets are indicated on the left and molecular weight markers on the right. Star indicates an irrelevant cross-reacting band seen with rabbit anti-HA.

**Figure 6.**
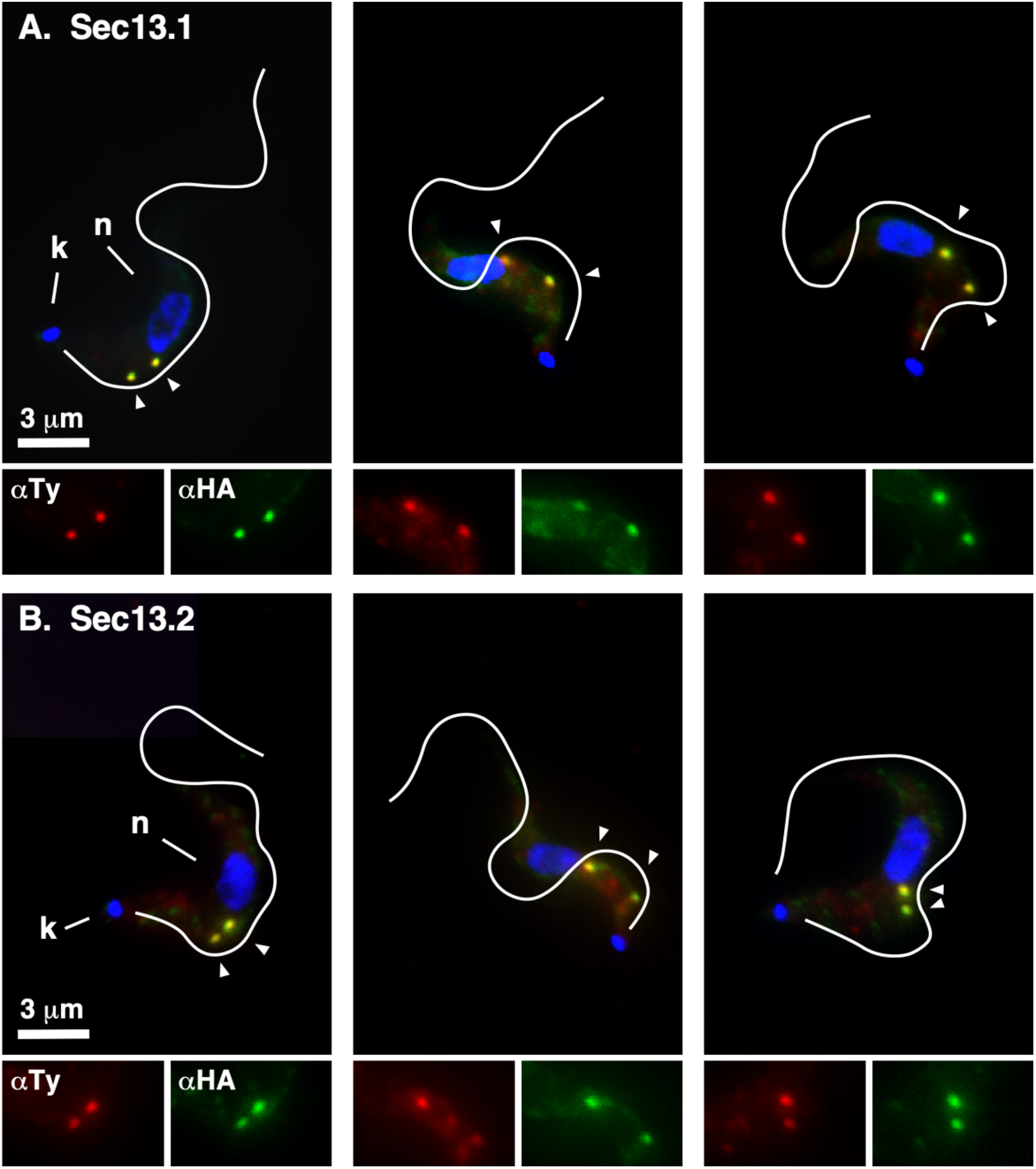
TbSec13 Localizations. Interphase BSF cells containing the Ty-tagged allele of *Sec24.1* as an ERES marker and HA-tagged alleles of *TbSec13.1* (A, top) or *TbSec13.2* (B, bottom) were stained with mAb anti-Ty (TbSec24.1, red) and rabbit anti-HA (TbSec13, green). Deconvolved summed-stack projections of individual cells are presented. White lines indicate the position of the flagellum as drawn from matched DIC images. Kinetoplasts (k) and nuclei (n) are indicated (left panels only). ERES are indicated by arrowheads. Enlarged single channel images of the ERES region are presented at the bottom of each image.

### Proximity Labeling of the ERES

To look closer at the presence of both TbSec13 orthologues in COPII structures, in particular TbSec13.2, we performed proximity labeling with the enhanced biotin ligase, TurboID [29]. An HA-tagged TurboID domain was fused *in situ* to the C-terminus of TbSec23.1 (TbSec23.1::Turbo::HA) in a parental BSF cell line bearing an *in situ* Ty-tagged TbSec24.2 orthologue (TbSec24.2::Ty) as an ERES marker [9]. Immunofluorescence analyses revealed that the Turbo-tagged TbSec23.1 colocalized precisely with TbSec24.2::Ty in the typical two ERESs of interphase cells (Fig. 7A, left). Similarly, streptavidin staining precisely overlapped with TbSec24.2::Ty indicating that the ERES is the predominant site of proximity biotinylation by TbSec23.1::Turbo::HA (Fig. 7A, right).

**Figure 7.**
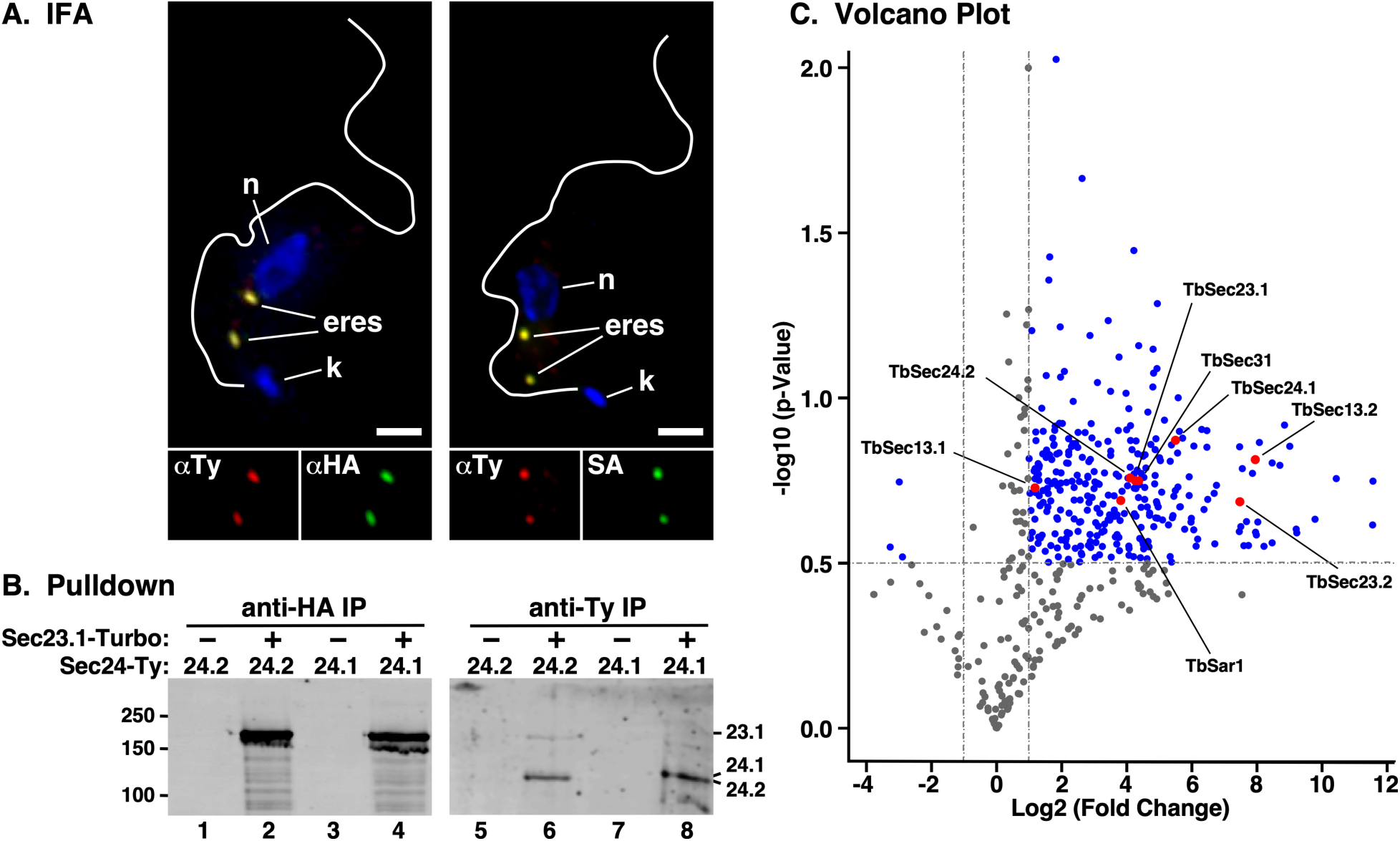
Proximity Labeling of the ERES. **A.** BSF cells co-expressing TbSec24.2::Ty and TbSec23.1::Turbo::HA were stained with anti-Ty (red) and anti-HA (green) (left panel) or with anti-Ty (red) and streptavidin (SA, green) (right panel). Cells were stained with DAPI to identify nuclei (n) and kinetoplasts (k). Deconvolved 3-channel summed stack projections are presented and ERES are indicated. Flagella outlines are from matched DIC images. Single channel images of the ERES region are shown (bottom). Bar, 2 μm. **B.** Lysates of TbSec24.1::Ty or TbSec24.2::Ty BSF cells alone (-) or co-expressing TbSec21.1::Turbo::HA (+) were subject to immunoprecipitation with anti-HA or anti-Ty antibodies as indicated. Pulldowns were fractionated by SDS-PAGE and affinity-blotted with streptavidin. Because the signal for TbSec23.1::HA was so intense the image was separated digitally and contrast enhanced independently for presentation. **C.** Volcano plot displaying protein hits from LC-tandem MS analysis (n=3) of PCF cells with (right side) and without (left side) expression of TbSec23.1::Turbo::HA. Log2 (Fold Change) and -log10(p-values) between the two conditions were calculated using R Studio. Dotted lines represent cutoff thresholds of -1 and 1 (vertical lines) for fold change and 0.5 (horizontal line) for statistical significance (p<0.05). Blue dots indicate significant protein hits, while red dots highlight the COPII subunits.

The functionality of the TbSec23.1::Turbo::HA reporter was confirmed by pull down assays. TbSec23.1::Turbo::HA was expressed in parental BSF cells bearing either TbSec24.1::Ty or TbSec24.2::Ty and immunoprecipitation from whole cell lysates was performed. The fractionated precipitates were then blotted with streptavidin. The parental TbSec24::Ty cell lines served as negative controls. As expected, strong auto-biotinylation was detected in the TbSec23.1::Turbo::HA cell lines following anti-HA pull down (Fig. 7B, lanes 2 & 4). No biotinylation was detected in the parental control cells (Fig. 7B, lanes 1 & 3) indicating strict dependence on the Turbo tagged reporter. In the anti-Ty pull downs, both biotinylated TbSec24.1::Ty and TbSec24.2::Ty were readily detected (Fig 7B, lanes 6 & 8), again dependent on the presence of the TbSec23 Turbo-tagged reporter (Fig 7B, lanes 5 & 7). Collectively, these data validate the proper localization of the TbSec23.1::Turbo::HA reporter, and its utility for proximity labeling of *bona fide* ERES components.

Biotinylated proteins were affinity purified from parental and TbSec23.1::Turbo::HA procyclic form (PCF) cell lines and subjected to LC-tandem-MS-based proteomic analyses (Fig. 7C). All of the COPII coat proteins and the COPII regulatory small GTPase Sar1 demonstrated statistically significant increased detection relative to parental cells, consistent with close proximity to the ERES, including TbSec13.2. Of these, TbSec13.1 demonstrated the smallest fold increase, likely because a significant portion of this protein is sequestered in nuclear pore complexes [20]. In contrast, TbSec13.2 showed the highest fold increase of any COPII component. While these results alone do not prove the presence of TbSec13.2 in COPII complexes, in conjunction with the localization and functional knockdown data, they are strongly supportive of this overall conclusion.

## DISCUSSION

The outer layer of the eukaryotic COPII machinery is comprised of Sec13/Sec31 heterotetramers that form a cage-like structure [16]. The single TbSec31 orthologue has been localized to the ERES in PCF trypanosomes, and RNAi knockdown indicates that it is an essential protein [30]. Likewise in PCF cells, TbSec13.1 localizes to the ERES [23]. Proteomic studies have also shown TbSec13.1 to be a *bona fide* component of the PCF nuclear pore complex (NPC), as it is in other systems, and that it localizes to puncta in the nuclear envelop [20, 21]. Although not commented on by these authors, their TbSec13.1 image ([20], Fig. 2A therein) also showed a prominent non-nuclear spot that is likely the nearby ERES. This dual localization is also noted in the TrypTag database [31], although it was annotated incorrectly as the adjacent Golgi, not the ERES. Recently, studies in the distantly related marine diplonemid *P. papillatum* found that the TbSec13.1 orthologue (PpSec13a) also localized to both NPC and ERES [24]. In regard to TbSec13.2, the combined proteomic [20, 21] and TrypTag data [31] are consistent with ERES localization, but not the NPC, suggesting a role in ER exit. In contrast, it was suggested that the *P. papillatum* orthologue (PpSec13b) is not associated with the ERES at all, based on negative proteomic data and failure to colocalize with PpSec13a, and consequently has no role in secretory trafficking [24]. Rather it was argued that PpSec13b is exclusively associated with the SEA/GATOR complex, and thus is likely involved in regulation of nutrient acquisition in the endolysosomal system. By analogy, this conclusion was extended to TbSec13.2.

Our imaging studies show clear association of TbSec13.1 with the ERES in BSF *T. brucei*, but little evidence of NPC localization. However, given the precedent for NPC association in multiple systems, we feel this likely represents differences in our tagging methodology and/or a lack of sensitivity. In this regard it has been shown recently that the NPC is less accessible to antibodies relative to smaller probes such as streptavidin [32]. Thus there is general agreement that the TbSec13.1 orthologue is involved in both secretory and nuclear transport processes, and our functional knock down studies support this conclusion in regard to secretion (discussed below). Likewise, we provide definitive colocalization evidence for TbSec13.2 in the ERES (with TbSec24.1 as the marker), consistent with the TrypTag assignment. This localization is strongly supported by our proximity labeling results, in which TbSec13.2 was an exceptionally robust hit, and by our functional knockdown studies (discussed below). In contrast, we found little evidence of TbSec13.2 in other post-nuclear (endolysosomal) localizations that would be consistent with the SEA/GATOR complex. It must also be noted that TrypTag did not assign additional endosomal localization to TbSec13.2 as was stated in [24]. Nevertheless, we do not consider our negative results sufficient to rule out such a function in *T. brucei*, in particular given that Sec13 orthologues are proven components of SEA/GATOR complexes in other systems, e.g., yeast and mammals [22]. However, our TbSec13.2 results contrast markedly with the *P. papillatum* orthologue (PpSec13b), for which no evidence of ERES localization was found [24]. It may well be that PpSec13b has been repurposed away from secretory trafficking in this distantly related euglenozoan, as suggested by these authors, but this is clearly not the case in *T. brucei*.

In all prior studies there was no direct assessment of the function of the two Sec13 orthologues in either *T. brucei* or *P. papillatum*. We have now performed detailed analyses of the roles of TbSec13.1 and TbSec13.2 in export of secretory cargo from the ER in BSF *T. brucei*. In considering these data it is worth noting that comparative proteomic analyses indicate that TbSec13.1 is ∼4-fold more abundant than TbSec13.2 in each life cycle stage, and that each protein has roughly similar abundance in BSF and PCF stages [33]. *<u>Firstly</u>*, we find that silencing of each orthologue is rapidly and selectively lethal in BSF trypanosomes, with TbSec13.1 being more sensitive (cessation of growth at 12 hrs vs. 24 hrs). *<u>Secondly</u>*, knockdown of each paralogue has largely similar effects on ER exit of secretory cargo: transport of GPI-anchored VSG was significantly reduced (3-4 fold); transport of soluble TbCatL was unaffected; and transport of transmembrane p67 was modestly impacted (1.8-2.8 fold). These effects are generally consistent with those we have seen previously with knock down of the inner TbSec23/24 COPII subunits [9]. These earlier studies also found no effect on transport of TbCatL, which is our most efficiently transported secretory reporter (lysosomal delivery *t*_1/2_ ∼10 min), and it is likely that this efficacy overrides the effects of TbSec13 knockdowns. In contrast, the large impact of each TbSec13 knockdown on transport of VSG, the overwhelmingly major secretory cargo of BSF trypanosomes, is consistent with the importance of efficiently synthesizing and transporting this protein to the cell surface [4]. Clearly, both trypanosomal Sec13 orthologues are required for this process. *<u>Thirdly</u>*, our prior work indicated that ER exit of VSG, and other GPI-anchored cargos, are selectively dependent on one of the two obligate Sec23:Sec24 heterodimers that form the inner layer of the COPII coat (Pair A: TbSec23.2:TbSec24.1) [9], and that this is mediated by transmembrane adaptors (TbERPs) that connect lumenal GPI-anchored cargo with the cytoplasmic COPII coat [14]. Our results here indicate that GPI selectivity is not influenced by TbSec13 orthologues, both of which must form heterotetramers with TbSec31 in the outer COPII coat. Overall then, the results of these trafficking assays fully confirm a COPII function for both TbSec13.1 and TbSec13.2 in the early secretory pathway in trypanosomes.

In summary, our findings inform a broader discussion of the diversification of components of the eukaryotic secretory machinery in the Euglenozoa, which include the sister groups kinetoplastids (*T. brucei*) and diplonemids (*P. papillatum*) [24]. Most members of these clades have two orthologues of Sec13, whereas other groups typically have single copies, e.g., vertebrates and fungi. This led Faktorova et al. to suggest that Sec13 gene duplication in the Euglenozoa has allowed a unique “division of labor” such that TbSec13.1 and PpSec13a function in nuclear and secretory transport processes, while TbSec13.2 and PpSec13b function in nutrient sensing via the SEA/GATOR complex, but not in secretory or nuclear transport.

However, our findings clearly demonstrate two overlapping sets of TbSec13 functions with the early secretory pathway being the common process, at least in the kinetoplastids. We would predict that this will be true in the diplonemids as well, but resolution of the issue will require direct functional experimentation. Fortunately, with the recent development of tools for genetic manipulation of *P. papillatum* this should be possible in the future [24].

## MATERIALS AND METHODS

### Maintenance of trypanosomes

All experiments (except proximity labeling) were performed in the single marker tetracycline-responsive Lister 427 strain *T. b. brucei* BSF cell line expressing VSG221 [34]. All cell lines were cultured in HMI-9 medium supplemented with 10% tetracycline-free fetal bovine serum (Atlanta Biologicals, Lawrenceville, GA) at 37°C in humidified 5% CO_2_ [35]. Cells were harvested at mid-to-late log phase (0.5-1×10^6^ cells/ml) for all experiments. For proximity labeling we used cultured procyclic form (PCF) cells of the Lister 427 strain [36]. Cell lines were cultured in Cunningham’s medium [37] supplemented with 10% tetracycline-free fetal bovine serum (Atlanta Biologicals, Lawrenceville, GA) at 27°C (site 2). All experiments were performed with cells harvested at the mid-to-late log phase (0.5-1×10^7^ cells/ml).

### Construction of RNAi, epitope-tagged, and proximity labeling cell lines

TbSec13.1 and TbSec13.2 RNAi constructs were generated in the pLEW100v5X:Pex11 stem-loop (pLEW100) vector [38]. The TbSec13.1 (nt 1-1110) and TbSec13.2 (nt 1-984) ORFs were PCR amplified from genomic DNA with flanking 5′ XhoI/XbaI and 3′ NdeI/AscI sites. The PCR products were sequentially inserted in one orientation downstream of the Pex11 stuffer using NdeI/XbaI and then upstream in the other orientation using XhoI/AscI. The resulting RNAi constructs were linearized with NotI and transfected independently into the single marker BSF cell line by electroporation [39] and clonal populations were selected on 24-well plates with phleomycin. dsRNA synthesis was induced with tetracycline (Tet: 1 μg/ml).

The generation and validation of the TbSec24.1::Ty and TbSec24.2::Ty *in situ* tagging constructs, and the generation of respective tagged BSF cell lines has been described previously [9]. *In situ* HA-tagged TbSec13.1::HA and TbSec13.2::HA were generated using the same methods. All three tagging constructs were liberated with KpnI/SacI and transfected into cultured BSF cells. First, we generated a clonal TbSec24.1::Ty cell line under neomycin selection. Expression of TbSec24 was confirmed by anti-Ty western blot. Next, this cell line was independently transfected with either the TbSec13.1::HA or TbSec13.2::HA construct and clonal double tagged cell lines were selected with neomycin/hygromycin. Expression of TbSec13.1::HA and TbSec13.2::HA positive cell lines were confirmed with Western blot (not shown).

To generate an ERES specific proximity labeling probe we first PCR amplified the C-terminus of the TbSec23.1 orf (nts 2238-2907) from genomic DNA and inserted it into the ClaI-HindIII sites of pXS6^(pur)^:3xHA upstream of the 3xHA tag [38]. Next the TurboID orf was PCR amplified from plasmid V5-TurboID-NES_pCDNA3 (a generous gift of Dr. Chris de Graffenried, Brown University) and inserted into EcoRI-XhoI sites between the Sec23.1 orf and the 3xHA tag creating an in frame fusion of TbSec23.1::TurboID::3xHA. Finally the TbSec23.1 3’ UTR (nts 1-653 relative to the stop codon) was PCR amplified from genomic DNA and inserted into PacI-SacI sites downstream of the puromycin resistance cassette. For purposes of validation the entire TbSec21.1::Turbo::HA construct [5’-3’: TbSec23.1::TurboID::3xHA / Aldolase IGR / Puromycin / TbSec23.1 3’ UTR] was excised with ClaI-SacI and electroporated into the TbSec24.1::Ty and TbSec24.2::Ty BSF cell lines described above. Clonal cell lines were obtained under puromycin selection and dual expression of Ty- and HA-tags was confirmed by Western blot and IFA (not shown). For large scale proximity labeling, an equivalent TbSec21.1::Turbo::HA *in situ* tagged PCF cell line was prepared and validated in the same manner.

### RNA extraction and qRT-PCR

Transcript levels of endogenous TbSec13 subunit genes were determined using quantitative reverse transcription PCR (qRT-PCR). Total RNA was isolated using RNeasy mini kit (Qiagen, Valencia, CA, USA). RNA was treated on-column with RNase-Free DNase (Qiagen, Valencia, CA, USA), and cDNA was prepared using iScript cDNA synthesis kit (Bio-Rad, Hercules, CA, USA) per manufacturer’s instructions. qRT-PCR reactions were prepared using Power SYBR Green PCR Master Mix (Life Technologies, Carlsbad, CA, USA), diluted cDNAs, and specific primers targeting the 3’ UTR region of endogenous TbSec13.1 (FP: 5’-GGGAAATGAGGACTATGGGAAG-3’ and RP: 5’-AAACTAGGAGGGTGAACTGTG-3’) or TbSec13.2 (FP: 5’-GGTAATACCGTCTGCTTGTAGG-3’ and RP: 5’-GAGGGATGCCAAACCAAGA-3’). The qRT-PCR reactions were performed in the StepOne™ Real-Time PCR System (Life Technologies, Carlsbad, CA, USA). Each reaction was performed in triplicates, and for each transcript, melting curves indicated a single dominant product post-amplification. Experimental transcripts were independently normalized to the internal reference gene TbZFP3 [40] Three biological replicates were performed for each TbSec23/24 subunit and means ± SD were quantified.

### Antibody, secondary, and blotting reagents

Rabbit anti-VSG117, rabbit anti-TbCatL, rabbit anti-BiP, and mouse monoclonal anti-p67 were described previously [28, 41, 42]. Mouse monoclonal anti-Ty ascites, and affinity purified rabbit anti-HA were generated by Convance Laboratories Inc. (Denver, PA, USA). Secondary reagents for IFA were A594 goat anti-mouse IgG, A488 goat anti-rabbit IgG, and A488 streptavidin (Molecular Probes, Eugene OR). IRDYe-800cw-streptavidin was used for blotting (LI-COR Biotech, Lincoln NE).

### Pulse/Chase Transport Assays

Pulse/chase metabolic radiolabeling with [^35^S]methionine/cysteine (Perkin Elmer, Waltham, MA, USA) and subsequent immunoprecipitation of radiolabeled proteins (VSG, TbCatL, and p67) from lysates and media fractions were performed as previously described with minor alterations [27]. In short, log phase cells were harvested, washed with Hepes-buffered saline (HBS: 50 mM HepesKOH, pH 7.5, 50 mM NaCl, 5 mM KCl, 70 mM glucose), and resuspended in methionine/cysteine-minus labeling media (10^8^/ml, 15 min, 27°C). Labeling was initiated by addition of [^35^S]Methionine/Cysteine (200 µC/ml, PerkinElmer, Waltham, MA); pulse times were 15 mins for VSG, 10 mins for TbCatL, and 15 mins for p67. The chase period was initiated by 10-fold dilution with prewarmed complete HMI9 medium, and samples (1.0 ml) were collected at specific time points as indicated in the relevant figures. For assay of TbCatL and p67 transport, sampled cells were washed with ice-cold HBS and solubilized in radioimmunoprecipitation assay buffer (RIPA: 50 mM Tris-HCl, pH 8.0, 150 mM NaCl, 1.0% NP-40, 0.5% deoxycholate, and 0.1% SDS) . Immunoprecipitated proteins were analyzed by SDS-PAGE and phosphorimaging using a Typhoon FLA 9000 with native ImageQuant Software (GE Healthcare, Piscataway, NJ, USA).

### Hypotonic Lysis Assay for VSG Transport

We used the established hypotonic lysis assay to determine transport of VSG to the cell surface [9, 11]. This assay relies on endogenous GPI-phospholipase C (GPI-PLC) to release surface VSG during hypotonic lysis. Internal VSG *en route* to the surface is resistant to this procedure and remains cell-associated. In brief, pelleted cells from the chase period samples were lysed with ice-cold dH_2_O (180 µl per sample; 10^6^ cells) with protease inhibitor cocktail (2 μg/ml each of leupeptin, antipain, pepstatin, and chymostatin) and Nα-tosyl-l-lysine chloromethyl ketone hydrochloride (TLCK; 0.37 μM/ml).

Samples were then supplemented with 20 μl of 10x TEN buffer (1x: 50 mM Tris-HCl, 150 mM NaCl, and 5 mM EDTA, pH 7.5) and incubated at 37°C for 10 mins to allow activated GPI-PLC to release soluble VSG from the cell surface. Time-dependent release during the pulse-chase corresponds to arrival at the cell surface. Cell and release fractions were separated by centrifugation, cells were solubilized in RIPA buffer, and supernatants were supplemented with RIPA detergents. Immunoprecipitation analyses of radiolabeled VSG polypeptides were performed as described above.

### Epifluorescence microscopy

Immunofluorescence (IFA) microscopy was performed as previously described [38, 43]. In short, log-phase BSF parasites were fixed with 2% formaldehyde and permeablized with 0.5% NP-40 followed by blocking, incubation with primary antibodies, and stained with appropriate Alexa488- or Alexa594-conjugated secondary antibodies. Slides were washed and mounted in DAPI fluoromount-G (Southern Biotech, Birmingham, AL) to reveal nuclei and kinetoplasts. Serial 0.2 micron image stacks (Z-increment) were collected with capture times from 100–500 msec (100x PlanApo, oil immersion, 1.46 numerical aperture) on a motorized Zeiss Axioimager M2 stand equipped with a rear-mounted excitation filter wheel, a triple pass (DAPI/FITC/Texas Red) emission cube, and differential interference contrast (DIC) optics. Images were captured with an Orca AG CCD camera (Hamamatsu, Bridgewater, NJ) in Volocity 6.0 acquisition software (Improvision, Lexington, MA), and individual channel stacks were deconvolved by a constrained iterative algorithm, pseudocolored, and merged using Volocity 6.1 Restoration Module. Images presented are summed stack projections of merged channels. The xyz pixel precision of this arrangement has been previously validated [9].

### BioID and MS Analysis

Initial validation experiments were performed in BSF cell lines containing the TbSec23.1::Turbo::HA *in situ* fusion and either the TbSec24.1::Ty or TbSec24.2::Ty *in situ* fusion. Because the TbSec23.1::Turbo::HA fusion is constitutively active, and trypanosomes require exogenous biotin for growth, proximity labeling is continuous during regular culture. Addition of exogenous biotin gave no additional benefit (not shown). Parental TbSec24.1::Ty or TbSec24.2::Ty (no Turbo) cell lines were used as controls. IFA and immunoprecipitation were performed as described above.

Proximity labeling for large scale affinity purification was done in the TbSec24.2::Ty PCF cell lines, without (control) or with (experimental) TbSec23.1::Turbo::HA. PCF cells grow to 10-fold higher density than BSF cells and each replicate (n=3) started with 1 L at 10^7^ cells/ml. Washed cells were lysed (10 ml at 10^9^ cells/ml) on ice in RIPA buffer with protease inhibitors [27]. Lysates were clarified by centrifugation and then rotated (4°C) overnight with 250 μl streptavidin beads (50% slurry, Sigma Aldrich, St Louis MO). Beads were washed 4x with RIPA buffer, and then 4x with 20 mM ammonium bicarbonate. Mass spectrometry was performed at the University at Buffalo Proteomic Core facility. Beads were subjected to a surfactant-aided precipitation and digestion protocol and eluted peptides processed for LC-MS analysis on an Ultimate 3000 nano-LC system coupled to an Orbitrap Fusion Lumos mass spectrometer.

Fractionated peptides were detected by a tandem scheme (MS1 Orbitrap; MS2 Ion Trap) in which the most abundant MS1 ions were selected, fragmented and acquired in MS2 scans to provide sequence specific information. Acquired spectra in each sample were matched to theoretical spectra generated from the TryTryp data base using Proteome Discover 1.4 (Thermo Fisher Scientific). Peptide-spectrum matches (PSM) were filtered and assembled to protein level by Scaffold 5 (Proteome Software, Portland OR), and protein/peptide false discovery rate (FDR) was controlled at 1% to ensure identification confidence. Data were subsequently processed as total ion chromatograph (TIC) counts. Experimental data sets (n=3) were aligned and hits not found in all sets were discarded, leaving 487 common proteins (Supplemental Table 1). The average TIC signal for each protein was calculated for each condition (control vs experimental). Log2 (Fold Change) and Log10 (p-value) were calculated and plotted in R software (https://www.r-project.org).

### Data Analyses

ImageJ (http://imagej.nih.gov/ij/) was used to quantify phosphorimages obtained from the Typhoon system. The intensities of specific bands (identical specific areas) within each lane were measured for quantification. To account for background noise, we independently subtracted the intensity of each specific band with an equivalent unlabeled area within the same lane. All subsequent data analysis was performed in Prism 9 (GraphPad Software Inc., San Diego, CA, USA).

## ACKNOWLEDGEMENTS

This work was supported by United States Public Health Service Grants NIAID R01 AI35739 to (JDB), and funds from the Jacobs School of Medicine and Biomedical Sciences (to JDB).

## SUPPLEMENTAL DATA

**Table S1. Protein List.** List of 487 common proteins identified by proximity labeling (TurboID) in all three replicates, ranked by fold change (Log2). Core COPII components are highlighted in yellow.

